# A novel olfactometer for efficient and flexible odorant delivery

**DOI:** 10.1101/461582

**Authors:** Shawn D. Burton, Mia Wipfel, Michael Guo, Thomas P. Eiting, Matt Wachowiak

**Author notes:** These authors contributed equally (Mia Wipfel, Michael Guo). Correspondence to be sent to: Matt Wachowiak, Department of Neurobiology and Anatomy, University of Utah, Salt Lake City, UT, USA.

## Abstract

Understanding how sensory space maps to neural activity in the olfactory system requires efficiently and flexibly delivering numerous odorants within single experimental preparations. Such delivery is difficult with current olfactometer designs, which typically include limited numbers of stimulus channels and are subject to inter-trial and inter-channel contamination of odorants. Here, we present a novel olfactometer design that is easily constructed, modular, and capable of delivering an unlimited number of odorants with temporal precision and no detectable inter-trial or inter-channel contamination. The olfactometer further allows for flexible generation of odorant mixtures and flexible timing of odorant sequences. Odorant delivery from the olfactometer is turbulent but reliable from trial-to-trial, supporting operant conditioning of mice in an odorant discrimination task and permitting odorants and concentrations to be mapped to neural activity with a level of precision equivalent to that obtained with a flow dilution olfactometer. This novel design thus provides several unique advantages for interrogating olfactory perception and for mapping sensory space to neural activity in the olfactory system.

## INTRODUCTION

Mapping sensory information to neural activity in the olfactory system is complicated by many factors, including the high dimensionality of olfactory stimulus space and limited knowledge of odorant receptor tuning. Overcoming these difficulties often requires testing large numbers of odorants in order to adequately cover even a portion of olfactory stimulus space, creating a need for methods of odorant delivery that are both efficient (i.e., tens-to-hundreds of odorants testable per preparation) and flexible (i.e., odorants chosen ‘on the fly’ during the course of an experiment).

Efficient and flexible odorant delivery is possible by manually presenting odorant reservoirs to the experimental preparation (e.g. see: (Takahashi et al., 2004)). However, such manual presentation involves long, variable, and relatively uncontrolled odorant delivery. Flow dilution olfactometers, which are widely used throughout olfactory research, instead enable automated delivery of temporally controlled streams of precisely diluted odorants using a combination of tubing, computer-controlled valves, and mass flow controllers (Bodyak and Slotnick, 1999; Slotnick and Restrepo, 2005; Johnson and Sobel, 2007; Schmidt and Cain, 2010). Flow dilution olfactometers have substantial limitations, however. For example, inter-trial and inter-channel contamination can arise from adsorption of odorants to surfaces downstream of odorant reservoirs (e.g., tubing, manifolds, mixing chambers, and final delivery ports), as well as from pressure fluctuations during valve actuation that can generate backflow of odorant vapor into upstream manifolds. Consequently, delivering numerous odorants with flow dilution olfactometers requires frequent and careful washing and/or replacement of multiple components, substantially reducing delivery efficiency and flexibility. Less standard flow dilution olfactometers employing large multi-channel odorant banks (e.g., see: (Davison and Katz, 2007; Soucy et al., 2009; Tan et al., 2010)) can achieve efficient odorant delivery, but are still subject to inter-trial and inter-channel contamination, provide limited delivery flexibility, and can be expensive to build.

Here, we describe and characterize an olfactometer capable of efficient and flexible odorant delivery with minimal inter-channel and inter-trial contamination. The novel design incorporates interchangeable modules with inexpensive and disposable odorant reservoirs that can be prepared during the course of an experiment, and uses spans of open airflow instead of tubing to eliminate common adsorptive surfaces that can give rise to contamination. Through these design features, the olfactometer generates turbulent yet temporally precise odorant delivery at adjustable concentrations, and further allows for flexible generation of vapor-phase odorant mixtures as well as ‘rapid-fire’ sequences of odorants. Employing this olfactometer to monitor sensory-evoked neural activity in the mouse olfactory system, we routinely test >100 odorants chosen ‘on the fly’ in a single experimental preparation, with multiple trials per odorant, and observe reliable odorant-evoked activity patterns with no detectable inter-trial or inter-channel contamination. We further demonstrate that the olfactometer readily supports behavioral testing in an operant conditioning odorant discrimination task. The novel olfactometer design thus provides several key advantages for probing olfactory perception and for mapping high-dimensional olfactory stimulus space to neural activity.

## MATERIALS AND METHODS

### Olfactometry

Odorants were acquired from Sigma-Aldrich, except for isoamyl acetate, which was acquired from MP Biomedicals, and methyl tiglate, which was acquired from Tokyo Chemical Industry Co., Ltd. Odorants were diluted in caprylic/capric medium chain triglycerides (C3465, Spectrum Chemical Mfg. Corp.). Characterization of olfactometer performance was completed using a photo-ionization detector (PID) (200B miniPID, Aurora Scientific), with the following olfactometer operating parameters held constant (unless otherwise stated): olfactometer-to-PID distance: 8-10 cm; channel pressure: 30 kPa; carrier stream flow rate: 8 L/min; odorant: (+)-α-pinene (1:5-10 liquid dilution). When evaluating the influence of each operating parameter on odorant delivery from the olfactometer, the parameter was systematically varied across interleaved trials (e.g., trial 1: 5 kPa channel pressure; trial 2: 10 kPa channel pressure, trial 3: 15 kPa channel pressure; etc.) using a single odorant reservoir to minimize any variability due to odorant depletion or slight differences in odorant reservoir preparation. To estimate the concentration of odorant delivery as a fraction of saturated odorant vapor, odorant delivery from the novel olfactometer and a previously described flow dilution olfactometer (Verhagen et al., 2007) were compared.

### Animals

For neural imaging experiments, olfactory marker protein (OMP)-Cre mice (Li et al., 2004) were crossed to Rosa26-CAG-lox-STOP-lox-GCaMP6f (RCL-GaMP6f) mice (Madisen et al., 2015). Female offspring heterozygous (**Figure 10**) or homozygous (**Figure 11**) for the OMP-Cre allele and 2-4 months in age were used. Mice were housed 5 per cage with food and water available *ad libitum*. For behavioral experiments, 3 C57BL/6 mice (1 male, 2 female) 2-6 months in age were used. All mice were kept on a 12 h light/dark cycle. All procedures were performed following the National Institutes of Health *Guide for the Care and Use of Laboratory Animals* and were approved by the University of Utah Institutional Animal Care and Use Committee.

### Neural imaging

Imaging was performed essentially as described previously (Wachowiak and Cohen, 2001). Briefly, mice were initially anesthetized with intraperitoneal injection of pentobarbital (50 mg/kg) and subcutaneous injection of chlorprothixene (12.5 mg/kg). Subcutaneous injection of atropine (0.5 mg/kg) was additionally provided to minimize mucus secretions and maintain nasal patency. Heart rate and body temperature were maintained at 300-500 beats per minute and 37°C, respectively, throughout the experiment. A double tracheotomy was performed and inhalation artificially controlled (rates: 1-3 Hz) via the ascending tracheal tube (Eiting and Wachowiak, 2018). For imaging, mice were head-fixed and the bone over the dorsal main olfactory bulbs thinned. Anesthesia was maintained by ~0.5% isoflurane delivered in pure O_2_ to the descending tracheal tube. Widefield epifluorescence signals were acquired with a 4×, 0.28 N.A. or 5×, 0.25 N.A. objective (Olympus) at 256×256 pixel resolution and 25 Hz frame rate using a back-illuminated CCD camera (NeuroCCD-SM256; RedShirtImaging) and Neuroplex software, with illumination provided by a 470 nm LED (M470L2, Thorlabs) and GFP filter set (GFP-1828A-000, Semrock).

### Behavior

Behavioral testing was modified from previously published protocols (Verhagen et al., 2007; Wachowiak et al., 2013). Briefly, mice were anesthetized with isoflurane, and a custom aluminum headbar was glued and then cemented (Teets Denture Material, Cooralite Dental Mfg. Co.) to the skull. Mice were given carprofen (5 mg/kg) and enrofloxacin (5-10 mg/kg) for 2 days post-surgery and housed individually with food and water available *ad libitum.* Approximately 1 week after headbar implantation, mice were water-restricted to provide motivation for behavioral testing. Specifically, mice were limited to 1.0 mL of water per day, until their body weight fell to 80-85% of their pre-restriction weight. Once criterion weight was reached, mice were habituated for 2-3 days to the experimenter, the behavioral testing apparatus, and the behavioral tasks. The apparatus consisted of an air-floated Styrofoam ball and bar capable of fastening to the implanted headbar, enabling head-fixation of the mouse while otherwise permitting free movement. A lickspout dispelling fresh water was placed ~5 mm from the mouth of the mouse, with water delivery in ~12 μL increments triggered by a solenoid valve upon correct task response. The task consisted of a standard go/no-go olfactory discrimination task, with 4-s-long odorant presentations. Mice were trained to lick for water reward during the presentation of a conditioned stimulus (CS+) odorant, and to withhold licking during an unconditioned stimulus (CS–) odorant.

### Data Analysis

Odorant delivery latencies were calculated as the delay from valve actuation to the first deviation of the PID signal ≥3×standard deviations (SDs) from the baseline. Odorant delivery durations were calculated as the difference between the first and last deviations of the PID signal ≥3×SD from the baseline. In neural imaging experiments, glomeruli were identified by manually drawing regions of interest (ROIs) around glomerular-sized foci of activity. ∆F/F responses were calculated by first subtracting the mean signal over the 1 s preceding odorant delivery onset (i.e., the resting fluorescence) from the mean signal over the 2-3 s following odorant delivery onset, and then dividing by the resting fluorescence. ROI ∆F/F timecourses and correlation values were then derived using the mean ∆F/F across all pixels within an ROI. For display, odorant response maps were filtered with a Gaussian kernel (σ = 0.75 pixels), upsampled with bilinear interpolation to 512×512 pixel resolution, and pseudocolored for clarity. All shaded regions and error bars depict mean ± SD.

## RESULTS

To enable efficient and flexible delivery of odorants with controllable timing, adjustable concentration, and minimal inter-trial and inter-channel contamination, we designed a novel modular olfactometer that uses spans of open air flow instead of tubing to both minimize adsorptive surfaces and facilitate rapid exchange of inexpensive disposable odorant reservoirs.

### Construction and operation

Similar to the general operating principle of flow dilution olfactometers, the novel olfactometer uses computer-controlled valves to direct air into an odorant reservoir, which expels saturated odorant vapor into a diluting carrier stream that is then directed to the experimental preparation. Distinct from flow dilution olfactometers, however, the novel olfactometer integrates spans of open airflow both immediately upstream and downstream of the odorant reservoirs (**Figure 1A**). The complete open flow of air downstream of the odorant reservoirs (**Figure 1B-E**) dramatically reduces the available surfaces to which expelled odorants can adsorb, thus minimizing inter-trial contamination of one odorant with another. Moreover, the spans of open airflow upstream of the odorant reservoirs eliminate the possibility of odorant backflow into upstream components, minimizing the potential for inter-channel contamination. Another critical feature of the novel design is the use of small disposable odorant reservoirs attached in sets of 12 to modular barrels. This feature enables numerous 12-odorant panels to be flexibly loaded and rapidly exchanged during an experiment.

**Figure 1.**
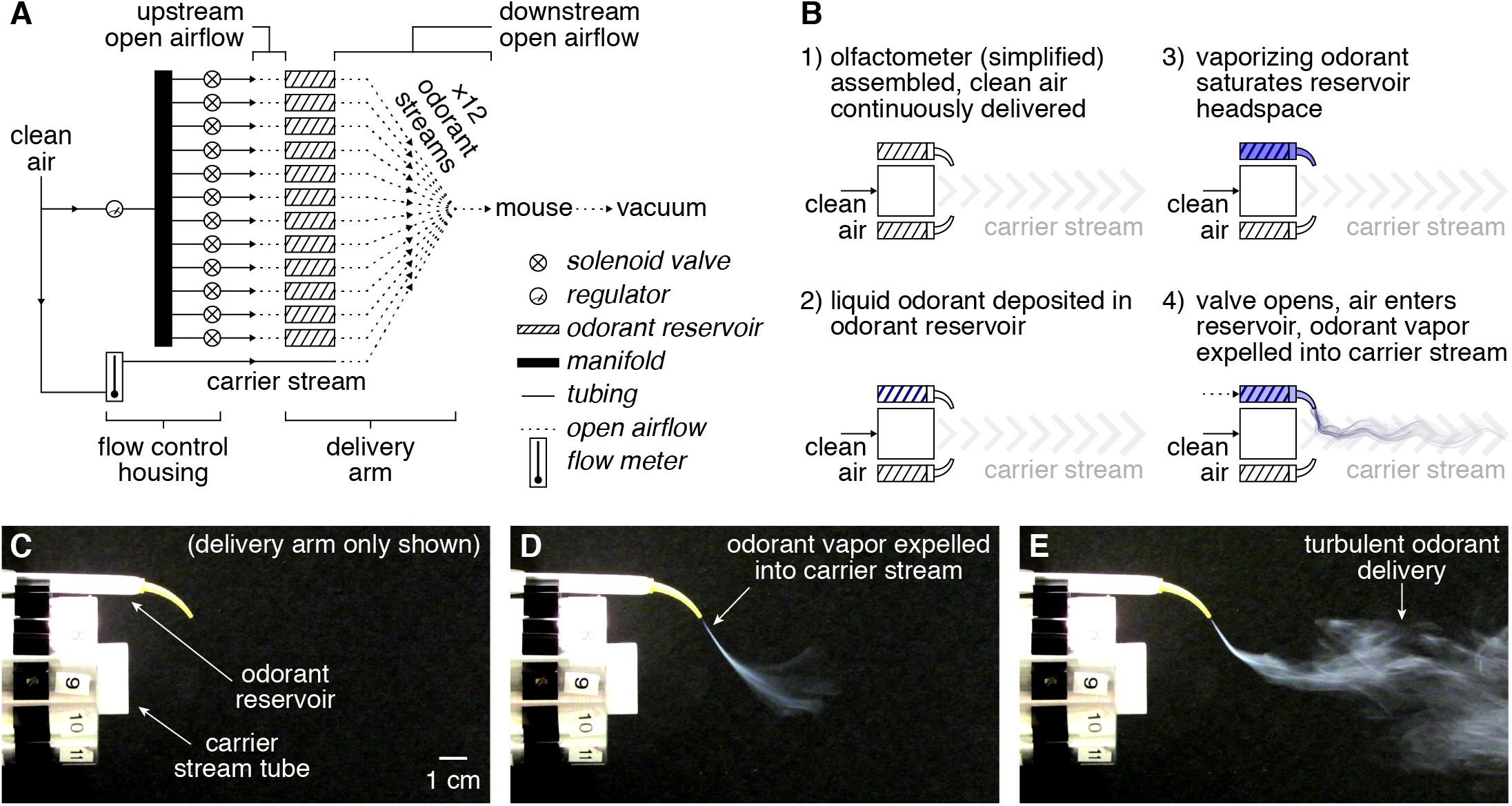
Operation of the novel olfactometer. (**A**) Pressurized clean air is directed through a manifold to 12 independent normally closed solenoid valves. Upon valve actuation, pressurized air flows across a short unconfined region (i.e., a span of open airflow) before entering a disposable odorant reservoir, expelling saturated odorant vapor from the reservoir into the central carrier stream. Bulk flow carries the final odorized stream across a span of open airflow (typically ~10 cm) to the experimental preparation (e.g., a mouse for neural imaging and/or behavioral testing). Odorized air not sampled by the preparation is continuously scavenged through an exhaust vacuum situated behind the experimental preparation. (**B**) Simplified illustration of operation, with two odorant reservoirs depicted for visual clarity. (**C-E**) Images of vaporized TiCl_4_ delivery from the olfactometer immediately before valve actuation (**C**), immediately following valve actuation (**D**), and following mixing of the expelled TiCl_4_ with the carrier stream (**E**). Only a single odorant reservoir (of 12 possible) was used for visual clarity.

The novel olfactometer includes two main components: 1) a delivery arm, which directs odorant delivery to the experimental preparation, and 2) a flow control housing, containing airflow controls (**Figures 1A, 2; Table 1**).

**Figure 2.**
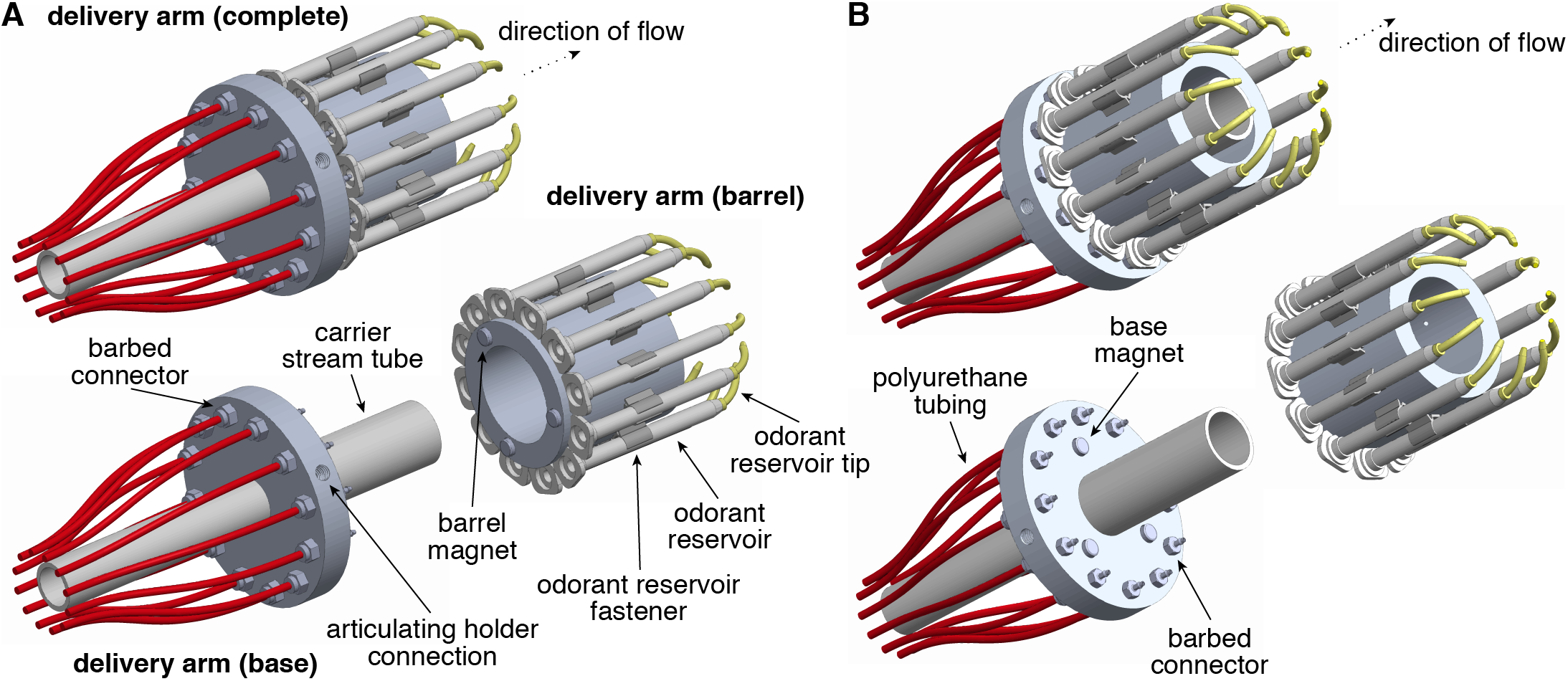
Construction of the novel olfactometer delivery arm. (**A,B**) Rotated view of the upstream (**A**) and downstream (**B**) surfaces of the novel olfactometer delivery arm, demonstrating magnetic attachment of modular barrels containing odorant reservoirs.

The delivery arm is comprised of a single base and multiple modular barrels (**Figure 2**). The base is constructed around a machined annular cylinder (O.D.: 8.60 cm [~3 ^3^/_8_ in]; I.D.: 1.91 cm [~^3^/_4_ in]; height: 1.27 cm [~^1^/_2_ in]) through which the carrier stream tube fits snugly. The carrier stream tube is further secured with a set screw (#10-32 thread) tapped into the base cylinder (not shown). In the design illustrated, we have used two concentric carrier stream tubes to broaden the final carrier stream of air at the delivery arm output, but in practice a single carrier stream tube suffices. An additional hole (#1/4-20 thread) tapped into the base cylinder is used to connect to an articulating dial gage holder (**Figure 2A**), allowing the delivery arm to be easily and stably positioned on any conventional air table. Twelve channels (#10-32 thread) are tapped through the base cylinder, and barbed connectors attached at both ends of each channel; connectors on the upstream surface (**Figure 2A**) connect to polyurethane tubing leading to independent solenoid valves within the flow control housing (**Figure 2B**), while connectors on the downstream surface (**Figure 2B**) direct airflow toward the bottom of each odorant reservoir when the barrel is assembled with the base. Importantly, these latter connectors do not physically touch the odorant reservoirs, creating a span of open airflow upstream of each reservoir (**Figure 1A**) that minimizes contamination and facilitates rapid exchange of pre-loaded barrels without requiring any components to be physically disconnected. To further accelerate this exchange and to ensure proper alignment of the base and barrel, three magnets are adhered in an asymmetric pattern to the downstream surface of the base cylinder (**Figure 2B**) and three more to the upstream surface of each barrel (**Figure 2A**).

The barrel is constructed around a machined annular cylinder (O.D.: 6.00 cm [~2 ^3^/_8_ in]; I.D.: 4.00 cm [~1 ^1^/_2_ in]; height: 6.00 cm [~2 ^3^/_8_ in]) that easily passes over the carrier stream tube (**Figure 2A,B**). Twelve holes (#2-56 thread) are tapped into the barrel cylinder (not shown) and used to attach the odorant reservoir fasteners (**Figure 2A**). Odorant reservoirs are repurposed mixers for two-part adhesives, with internal helical projections providing extensive surface area over which odorant solution can distribute to yield a disposable reservoir with small headspace that rapidly saturates with odorant vapor (**Figure 1B**). These repurposed mixers are mass-produced for a variety of applications, and so are relatively inexpensive and readily available in large quantities. Odorant reservoirs easily snap into the fasteners with slight pressure, and are capped with tightly fitting disposable curved tips (**Figure 2A**) that direct the expelled saturated odorant vapor from the reservoir into the central carrier stream. Odorant reservoir tips are repurposed dental applicators that, like the odorant reservoirs, are also relatively inexpensive and readily available in large quantities.

**Table 1.**
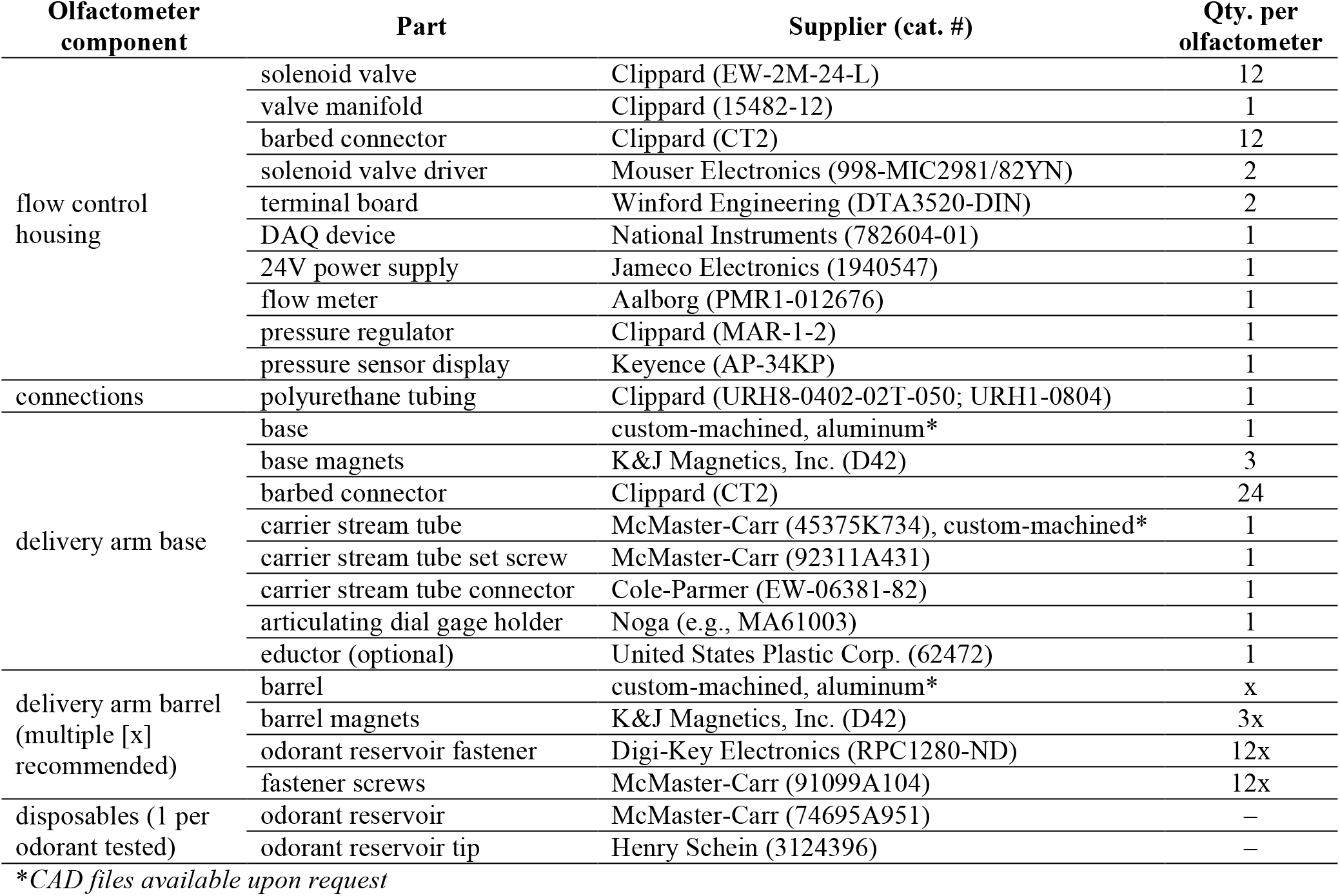
Olfactometer parts list.

The parameters of odorant delivery, including which of the odorants loaded into the modular barrel are delivered to the experimental preparation, as well as the specific time and concentration of odorant delivery, are governed by components within the flow control housing (**Figure 1A**). Clean air enters the flow control housing and is passed to the upstream end of the delivery arm carrier stream tube through a polyurethane tube at a constant rate controlled by a simple manual flow meter. In parallel, clean air from a separate source (or split from the same source as above) is split by a manifold into 12 separate channels, each of which connects to the inlet of an independently controlled, normally closed solenoid valve. A single miniature manual regulator upstream of the manifold controls the pressure of clean air at all 12 valves, such that changes in pressure uniformly influence delivery across all 12 channels. Polyurethane tubing connects valve outlets to the corresponding barbed connectors on the upstream surface of the delivery arm base. Odorant delivery is achieved and temporally controlled by actuation of a valve to the open state, which directs pressurized clean air toward the upstream end of an odorant reservoir, expelling saturated odorant vapor into the continuously flowing carrier stream approximately 5-10 cm from the downstream end of the carrier stream tube (**Figure 1B-E**). In our implementation, valve actuation is controlled by National Instruments DAQ hardware and custom National Instruments LabVIEW code, though we note that other, more economical alternatives (e.g., Arduino) could easily be employed. An exhaust vacuum to scavenge excess odorant not sampled by the experimental preparation (**Figure 1A**) is highly recommended, given the relatively broad stream of odorant vapor delivered by the olfactometer. Clearance of excess odorant is further promoted by the continuous delivery of clean air via the carrier stream when all 12 valves are closed, which additionally minimizes potential nonspecific odorant input to the experimental preparation arising from the laboratory environment or the experimental preparation itself.

At current prices, construction of the novel olfactometer (excluding the few custom-machined components) totals ~$1,500.00, with each odorant tested costing an additional ~$1.00 in disposables (i.e., odorant reservoir and odorant reservoir tip).

### Characterization of odorant delivery

Visualization of odorant delivery from the novel olfactometer using vaporized TiCl_4_ revealed turbulent plumes (**Figure 1E**). Effective implementation of the olfactometer will thus require careful consideration and understanding of this variable stimulus profile and its dependence on various operating parameters. We therefore next systematically characterized the performance of the olfactometer using a PID across a range of operating parameters.

**Figure 3.**
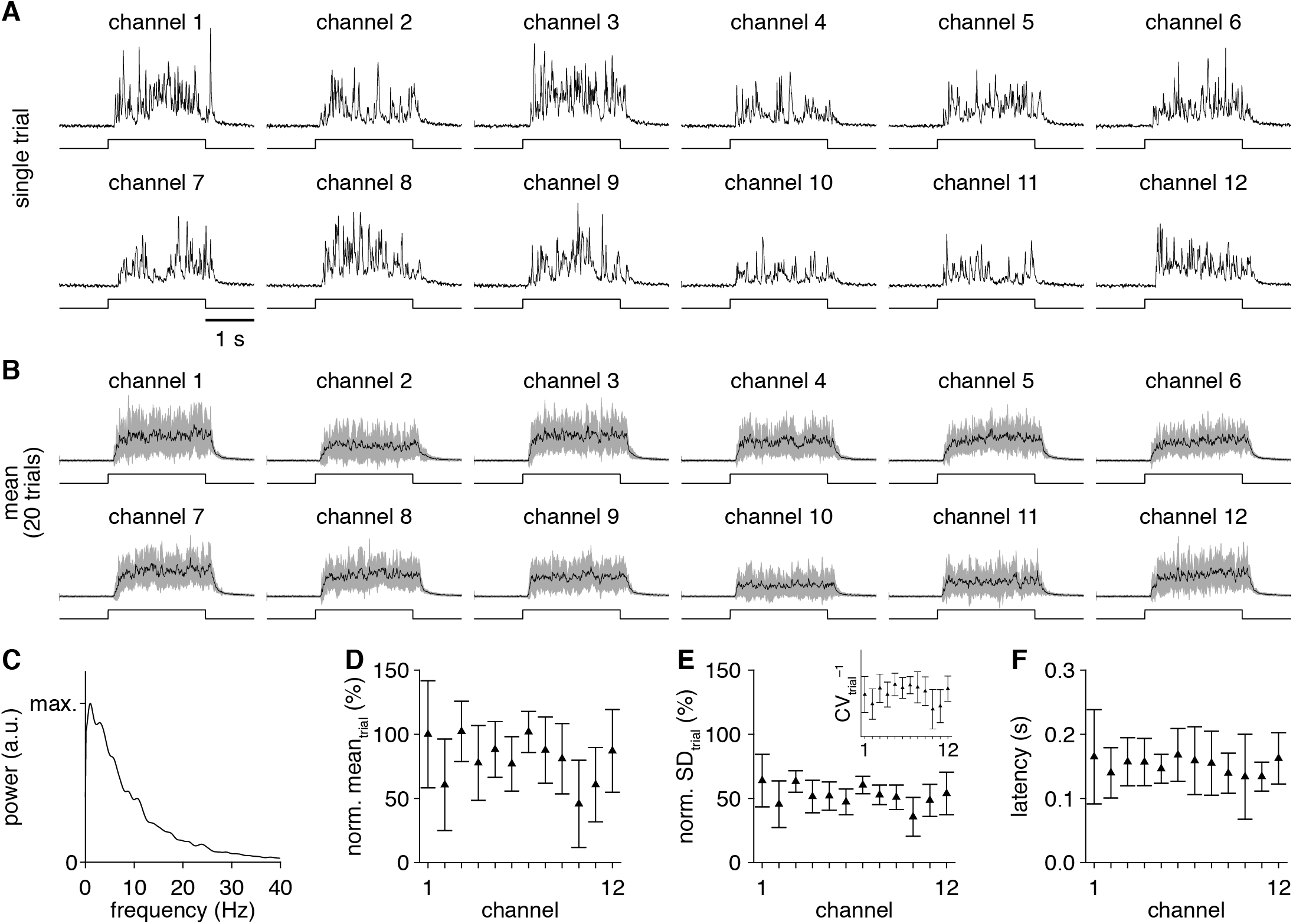
Comparable odorant delivery across 12 distinct channels. (**A**) PID recordings (upper) during single trials of 2-s-long valve openings (lower) from each olfactometer channel. (**B**) Mean PID recording across 20 trials for each olfactometer channel. (**C**) Mean power spectrum, averaged across all channels and trials. (**D,E**) PID recording mean (**D**) and SD (**E**) during the odorant delivery for each channel (normalized to the channel 1 mean). Inset: inverse of the coefficient of variation (CV) for each channel, reflecting the strength of odorant delivery relative to the delivery variability. (**F**) Mean latency of odorant delivery for each channel.

We first examined the profile and consistency of odorant delivery from the 12 olfactometer channels. On single trials, 2-s-long valve openings drove turbulent delivery of odorant (**Figure 3A**) with PID signals spanning frequencies of ~0-20 Hz (**Figure 3C**). While turbulent, the overall envelope of odorant delivery was reliable from trial-to-trial, such that averaging PID signals across multiple trials yielded step-like profiles (**Figure 3B**) comparable to the profile of odorant delivery from a flow dilution olfactometer. Delivery was similar across each of the 12 channels, with comparable odorant delivery mean (i.e., concentration) and variance (**Figure 3D,E**). Likewise, the latency from valve opening to odorant delivery was consistent across the 12 channels (**Figure 3F**), though longer than observed with flow dilution olfactometers, which are typically positioned much closer to the experimental preparation (i.e., 0-2 cm vs. 5-10 cm). Repositioning the PID to upstream of the odorant reservoir revealed no or negligible odorant backflow (data not shown), and in practice any such backflow would be scavenged by typical exhaust systems positioned around the experimental station. In total, the olfactometer thus mediates turbulent but overall reliable and comparable odorant delivery across 12 independent channels.

Saturated odorant vapor expelled from the odorant reservoir visibly disperses within the carrier stream as distance from the olfactometer increases (**Figure 1E**). Systematically varying the distance between the PID and the downstream end of the carrier stream tube quantitatively confirmed this observation (**Figure 4A,B**). The highest mean concentration of odorant was detected where the expelled saturated odorant vapor first entered the carrier stream (~6 cm), with monotonic decreases in delivery mean (**Figure 4C**) and variance (**Figure 4D**) and a monotonic increase in latency as distanced increased (**Figure 4E**). At the longest distances tested, the profile of odorant delivery further exhibited fewer high-frequency components (**Figure 4F**). Even with the distance-dependent decline in mean concentration, however, reliable odorant delivery with predictable latencies was still achieved with olfactometer-to-PID distances up to ~12 cm. The olfactometer can thus easily accommodate a variety of experimental paradigms (e.g., the concurrent positioning of a lickspout for operant conditioning).

**Figure 4.**
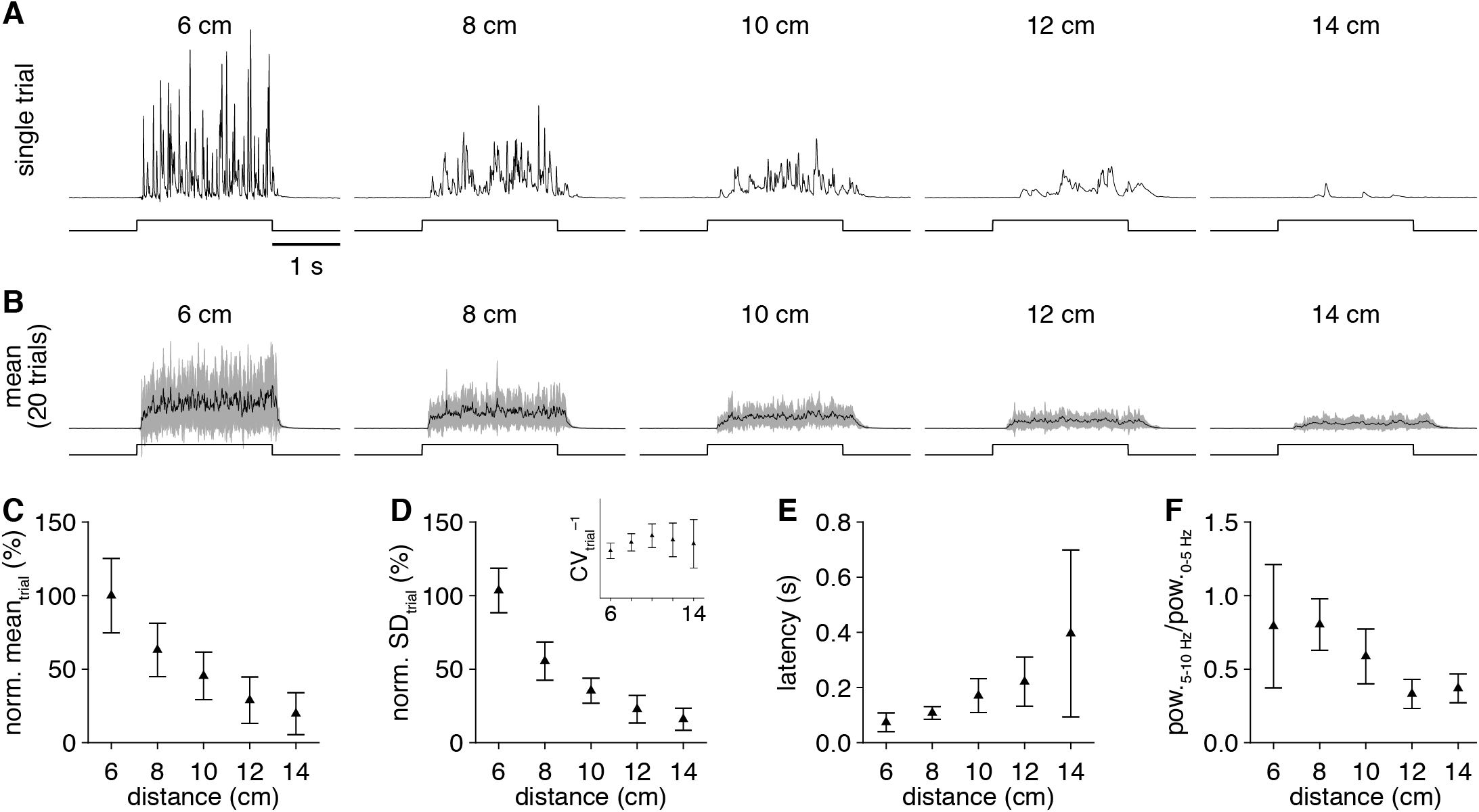
Varying the olfactometer-to-experimental preparation distance modulates the concentration of delivered odorant. (**A**) PID recordings (upper) during single trials of 2-s-long valve openings (lower) from a single olfactometer channel with varying olfactometer-to-PID distances. (**B**) Mean PID recording across 20 trials for each distance tested. (**C,D**) PID recording mean (**C**) and SD (**D**) during the odorant delivery for increasing distance (normalized to the 6 cm-distance mean). Inset: inverse of the CV for each distance. (**E**) Mean latency of odorant delivery for increasing distance. (**F**) Mean ratio of high frequency (5-10 Hz) to low frequency (0-5 Hz) spectral components in the PID recordings for increasing distance.

On first principles, increasing the pressure of clean air at the channel valve should increase the rate of airflow through the odorant reservoir upon valve actuation, thus increasing the amount of saturated odorant vapor expelled into the carrier stream. Indeed, we observed a monotonic increase in the mean concentration (and variance) of odorant delivery (**Figure 5A,B,D,E**) with increasing channel pressure. In contrast, there was no detectable change in the ratio of high-to-low frequency components in the odorant delivery profile (data not shown), and the latency of odorant delivery was minimally impacted at channel pressures ≥10 kPa and essentially identical for channel pressures 20-40 kPa (**Figure 5F**). Varying channel pressure thus provides a convenient avenue for instantaneously modulating the mean concentration of odorant delivery. To estimate the range of concentrations achievable through channel pressure modulation in an intuitive format, we directly compared delivery of the same odorant from the novel olfactometer vs. a flow dilution olfactometer (**Figure 5C;** see **Materials and Methods**). Relating the PID signals obtained with varying channel pressures vs. varying air dilutions of saturated vapor, we found that channel pressure can modulate the mean concentration of odorant delivery over a range equivalent to ~0.5-2.5% saturated odorant vapor (**Figure 5G**), depending on other operational parameters (e.g., decreasing the olfactometer-to-experimental preparation distance will shift this range to higher values).

**Figure 5.**
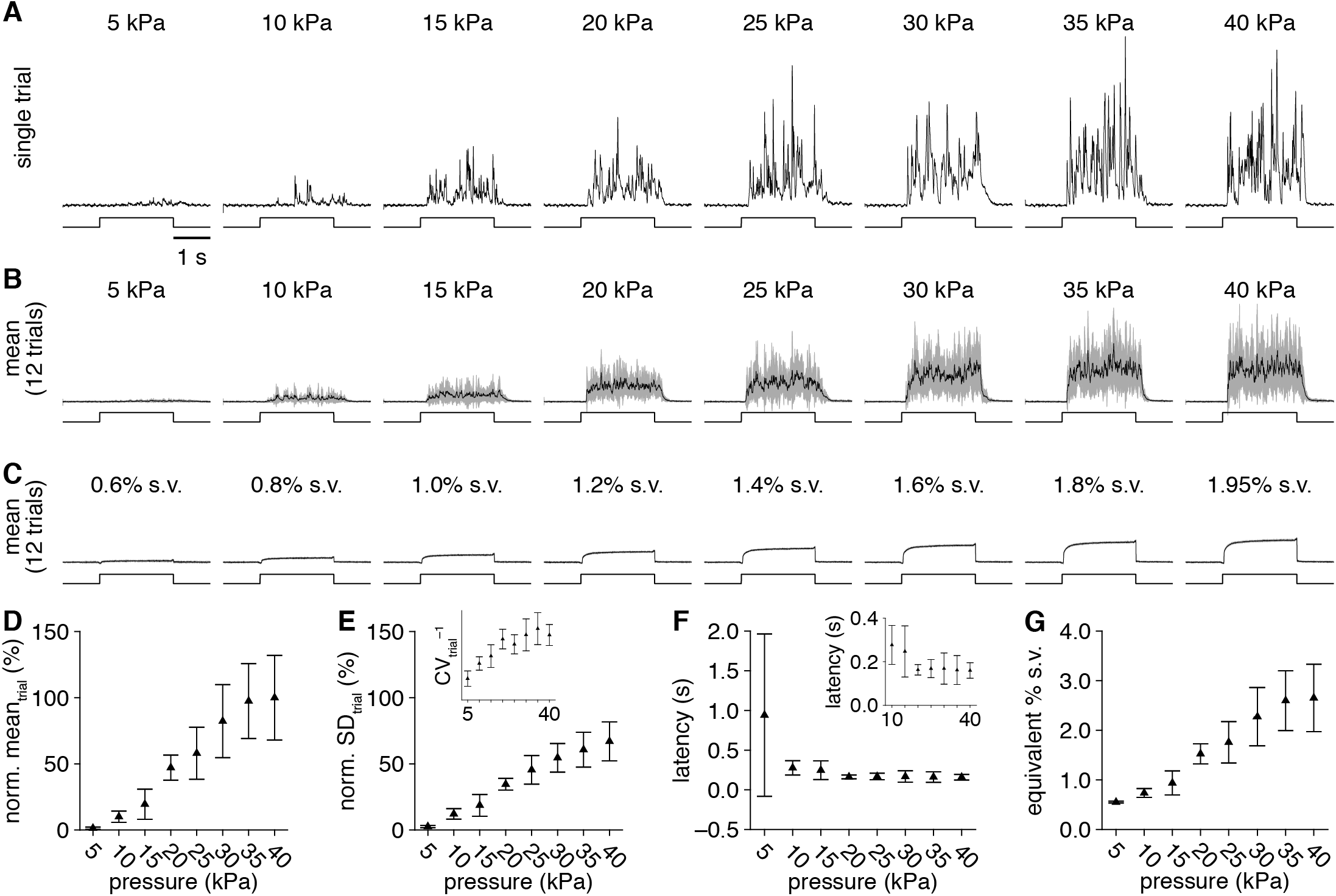
Varying the olfactometer channel pressure modulates the mean concentration of odorant delivery. (**A**) PID recordings (upper) during single trials of 2-s-long valve openings (lower) from a single olfactometer channel with varying channel pressure. (**B**) Mean PID recording across 12 trials for each channel pressure tested. (**C**) For comparison, mean PID recordings (upper) over 12 trials of 2-s odorant delivery (lower) of varying-percent saturated vapor from a flow dilution olfactometer. (**D,E**) PID recording mean (**D**) and SD (**E**) during the odorant delivery for increasing channel pressure (normalized to the 40 kPa-channel pressure mean). Inset: inverse of the CV for each channel pressure, demonstrating maximal odorant delivery with minimal variability at 30-40 kPa. (**F**) Mean latency of odorant delivery for increasing channel pressure. Inset: magnification of latencies for pressures ≥10 kPa. (**G**) Relationship between channel pressure and equivalent percent saturated vapor of odorant delivery, calculated from the PID recording mean obtained from the novel olfactometer and a flow dilution olfactometer.

In contrast to changes in channel pressure, varying the carrier stream flow rate strongly modulated the temporal properties of odorant delivery. Lower flow rates yielded both longer and more variable odorant delivery latencies (**Figure 6A-C**), as well as fewer high frequency components in the profile of odorant delivery (**Figure 6A,D**), consistent with previous results using a fan to generate turbulent odorant plumes (Nagel and Wilson, 2011). Surprisingly, with the exception of flow rates ≤2 L/min (which resulted in unreliable odorant delivery), this modulation of temporal properties occurred with minimal changes in the mean concentration of odorant delivery (**Figure 6B,E**), suggesting that the expelled saturated odorant vapor is incompletely mixed in the carrier stream. Modulating the carrier stream flow rate thus provides a simple method for investigating how varying plume dynamics impact neural activity and olfactory perception.

**Figure 6.**
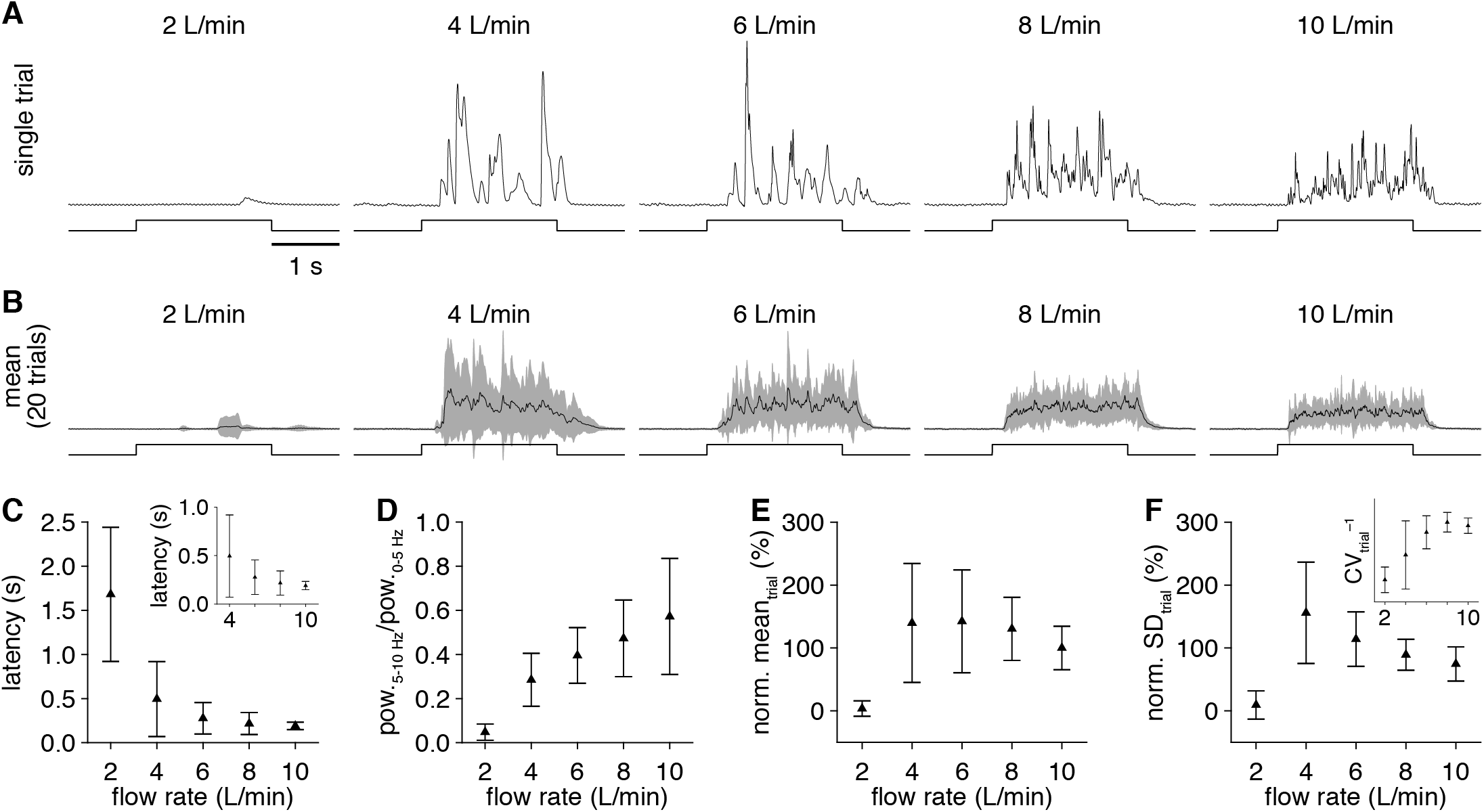
Varying the olfactometer carrier stream flow rate modulates odorant plume dynamics. (**A**) PID recordings (upper) during single trials of 2-s-long valve openings (lower) from a single olfactometer channel with varying carrier stream flow rate. (**B**) Mean PID recording across 20 trials for each flow rate tested. (**C**) Mean latency of odorant delivery for increasing flow rate. Inset: magnification of latencies for flow rates ≥4 L/min. (**D**) Mean ratio of high frequency (5-10 Hz) to low frequency (0-5 Hz) spectral components in the PID recordings for increasing flow rate. (**E,F**) PID recording mean (**E**) and SD (**F**) during the odorant delivery for increasing flow rate (normalized to the 10 L/min flow rate mean). Inset: inverse of the CV for each flow rate, demonstrating maximal odorant delivery with minimal variability at 8 L/min flow rate.

Different experimental paradigms require variable stimulus durations. We therefore next examined the resolution of odorant delivery duration afforded by the novel olfactometer. Valve opening durations as short as 20 ms yielded detectable odorant delivery on most trials (**Figure 7A,B**), with a minimal detected odorant delivery duration (see **Materials and Methods**) of 72.6±64.3 ms (**Figure 7C**), though this likely differs for varying operating parameters. Slightly longer valve opening durations (≥50 ms) yielded more reliable odorant delivery (**Figure 7D**), and all valve opening durations ≥100 ms delivered odorant at essentially identical rates (**Figure 7E**) and latencies (**Figure 7F**), reflecting the step-like mean odorant delivery profile. The novel olfactometer is thus capable of temporally precise odorant delivery across a wide range of stimulus durations.

**Figure 7.**
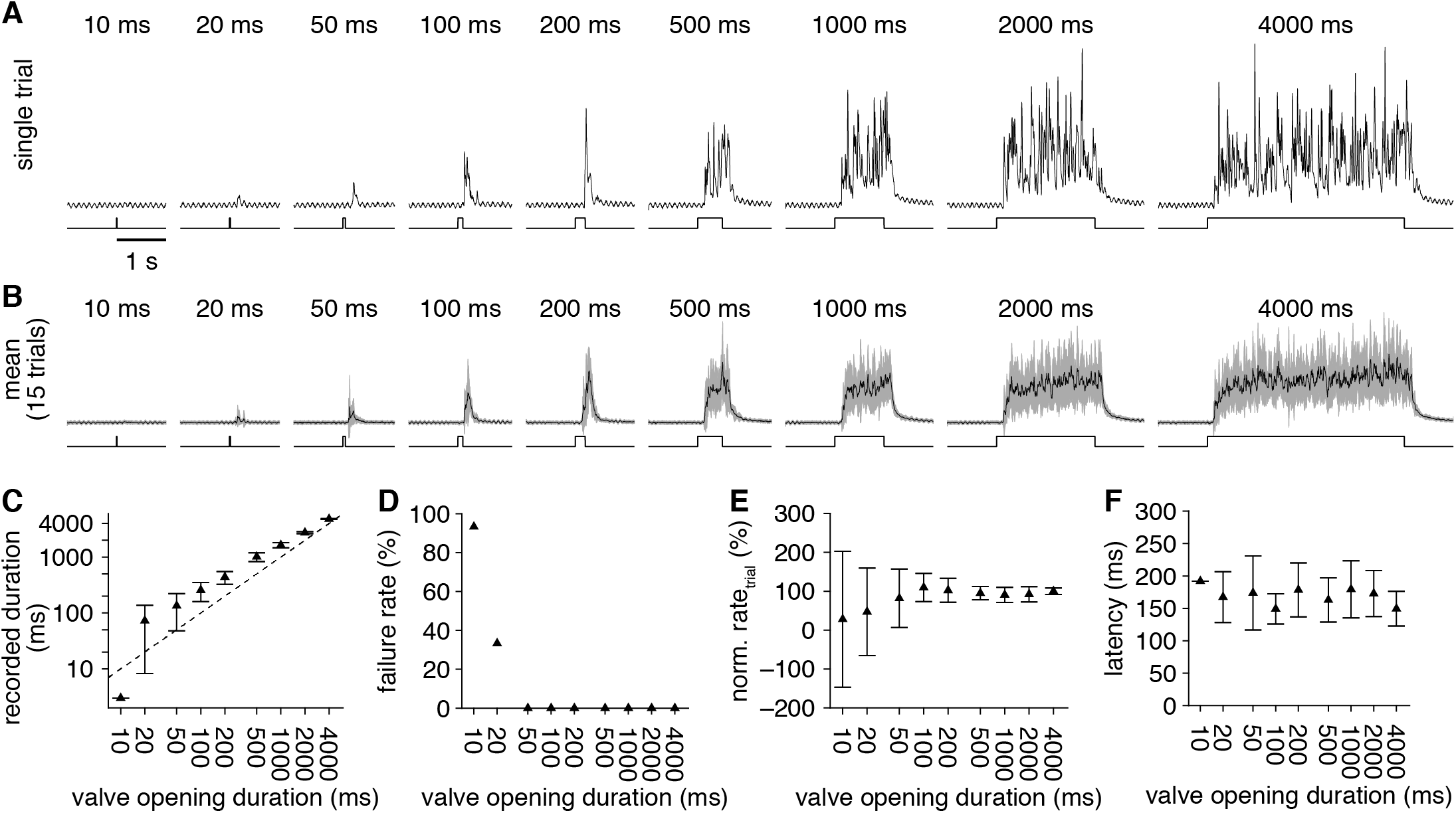
Tightly controlled odorant delivery duration. (**A**) PID recordings (upper) during single trials of odorant delivery with increasing valve opening duration (lower) from a single olfactometer channel. (**B**) Mean PID recording across 15 trials for each valve opening duration tested. (**C**) Mean duration of odorant delivery detected for increasing valve opening duration. Axes plotted on a log-scale. Dashed line: unity. (**D**) Mean odorant delivery failure rate (i.e., percent of trials with no detectable odorant-evoked PID signal) for increasing valve opening duration. (**E**) Mean odorant delivery rate, calculated as the PID recording mean divided by the valve opening duration (normalized to the mean delivery rate with 4000 ms-valve opening duration). (**F**) Mean latency of odorant delivery for increasing valve opening duration.

The novel olfactometer intentionally uses small odorant reservoirs for their disposability, enabling numerous odorants to be delivered without requiring careful maintenance of more permanent reservoirs (e.g., reservoir cleaning, capping loaded reservoirs with inert gas to limit oxidation, etc.). As a consequence of the necessarily small volumes of odorants used, however, odorant volatility may impact the stability of odorant delivery across time. Specifically, odorant delivery across several trials (or prolonged odorant delivery across fewer trials) may substantially deplete the small volume of high volatility odorants. To investigate this possibility, we examined the stability of odorant delivery across 50 consecutive trials of 2-s-long valve openings for 22 chemically diverse odorants (**Table 2**) ranging from low vapor pressure (**Figure 8A,B**) to high vapor pressure (**Figure 8C,D**). As expected, the concentration of odorant delivery decreased across trials in a vapor pressure-dependent manner (**Figure 8E**). While this decay is thus a limitation of the novel olfactometer design, we note that odorant depletion was slow (i.e., τ_delivery_>100 s) for the majority of odorants tested, and that even high volatility odorants (e.g., 2-butanone) yielded several seconds worth of relatively stable delivery sufficient for multiple trials of neural imaging and/or behavioral testing.

**Figure 8.**
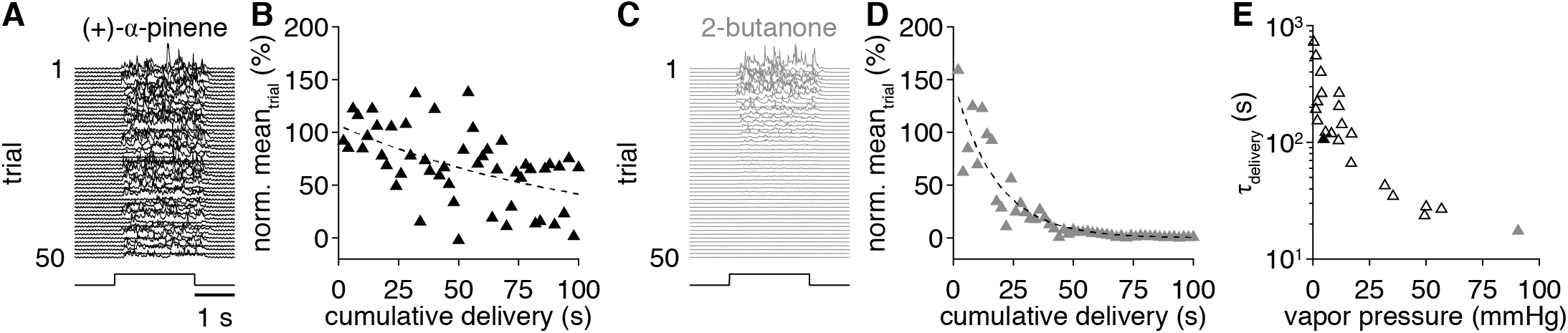
Odorant volatility influences the stability of odorant delivery. (**A**) PID recordings (upper) across 50 consecutive trials of 2-s delivery (lower) of a low volatility odorant: (+)-α-pinene (1:10 liquid dilution). (**B**) PID recording mean during the odorant delivery across 50 consecutive trials (100 cumulative seconds) from **A**. Dashed line: exponential fit. (**C,D**) Same as **A,B** for a high volatility odorant: 2-butanone (1:10 liquid dilution). (**E**) Odorant delivery decay (i.e., τ_delivery_; from single exponential fits, as in **B,D**) for 22 odorants with varying volatility. Solid black and grey points correspond to odorants shown in **A,B** and **C,D**, respectively.

**Table 2.** Vapor pressure-dependence of odorant delivery stability.

| Odorant* | CAS # | Vapor pressure<br>(mmHg, 25.0 C) | $\tau_{\text{delivery}}$<br>(s) |
| --- | --- | --- | --- |
| acetophenone | 98-86-2 | 0.397<sup>†</sup> | 721.09 |
| benzaldehyde | 100-52-7 | 1.27<sup>†</sup> | 191.79 |
| 2-octanone | 111-13-7 | 1.35<sup>†</sup> | 551.68 |
| R-(+)-limonene | 5989-27-5 | 1.98<sup>†</sup> | 222.73 |
| 4-methylanisole | 104-93-8 | 2.029<sup>‡</sup> | 153.18 |
| heptaldehyde | 111-71-7 | 3.52<sup>†</sup> | 398.78 |
| 2,6-dimethylpyrazine | 108-50-9 | 3.873<sup>‡</sup> | 262.36 |
| (+)-α-pinene | 7785-70-8 | 4.75<sup>†</sup> | 106.20 |
| isoamyl acetate | 123-92-2 | 5.6<sup>†</sup> | 122.00 |
| 4-methyl-3-penten-2-one | 141-79-7 | 8.21<sup>†</sup> | 118.77 |
| hexaldehyde | 66-25-1 | 11.3<sup>†</sup> | 103.50 |
| butyl acetate | 123-86-4 | 11.5<sup>†</sup> | 202.83 |
| 2-hexanone | 591-78-6 | 11.6<sup>†</sup> | 263.91 |
| ethyl butyrate | 105-54-4 | 12.8<sup>‡</sup> | 142.13 |
| 2-methylvaleraldehyde | 123-15-9 | 16.915<sup>‡</sup> | 66.26 |
| trans-2-methyl-2-butenal | 497-03-0 | 17.107<sup>‡</sup> | 117.96 |
| valeraldehyde | 110-62-3 | 31.792<sup>‡</sup> | 42.52 |
| 2-pentanone | 107-87-9 | 35.4<sup>†</sup> | 34.51 |
| 2-methylbutyraldehyde | 96-17-3 | 49.317<sup>‡</sup> | 23.58 |
| isovaleraldehyde | 590-86-3 | 50<sup>†</sup> | 28.05 |
| butane-2,3-dione | 431-03-8 | 56.8<sup>†</sup> | 26.77 |
| 2-butanone | 78-93-3 | 90.6<sup>†</sup> | 17.43 |

\*prepared at 1:10 liquid dilution
<sup>†</sup>PubChem (https://pubchem.ncbi.nlm.nih.gov/)
<sup>‡</sup>The Good Scents Company Information System (http://www.thegoodscentscompany.com/)

In principle, the novel olfactometer can flexibly generate any of 4083 distinct odorant mixtures per 12-odorant panel (i.e 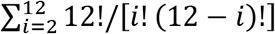) through the simultaneous actuation of multiple valves, with minimal concern over inter-channel contamination given the open airflow design. In practice, however, the dependence of odorant delivery concentration on channel pressure (**Figure 5**) together with the drop in channel pressure caused by each open valve will limit the total number of simultaneously active channels over which the olfactometer can achieve reliable odorant delivery. To determine this upper limit, we therefore next measured odorant delivery from a single channel while co-activating an increasing number of “blank” channels (i.e., channels with non-photoionizable odorant solvent only loaded into odorant reservoirs), using 40 kPa as a baseline channel pressure. As expected, co-activation of 1 (**Figure 9A,B**) or 3 (**Figure 9C,D**) blank channels reduced the mean concentration of odorant delivery (**Figure 9G**), though still yielded highly reliable odorant delivery with consistent trial-to-trial latencies (**Figure 9I**). Indeed, reliable odorant delivery could still be detected with co-activation of 7 blank channels (**Figure 9E,F**), albeit at much lower concentrations (**Figure 9G**) and slightly more variable latencies (**Figure 9I**). We thus take 8 as the maximum number of simultaneously active channels that the olfactometer can achieve reliable odorant delivery over (given a 40 kPa baseline channel pressure), supporting the flexible generation of any of 3784 possible odorant mixtures per trial. Depending on the maximal channel pressure afforded by the clean air source, this limit can also be readily increased to 12 simultaneously active channels and 4083 possible mixtures by increasing the baseline channel pressure to offset the division of airflow across simultaneously open valves.

**Figure 9.**
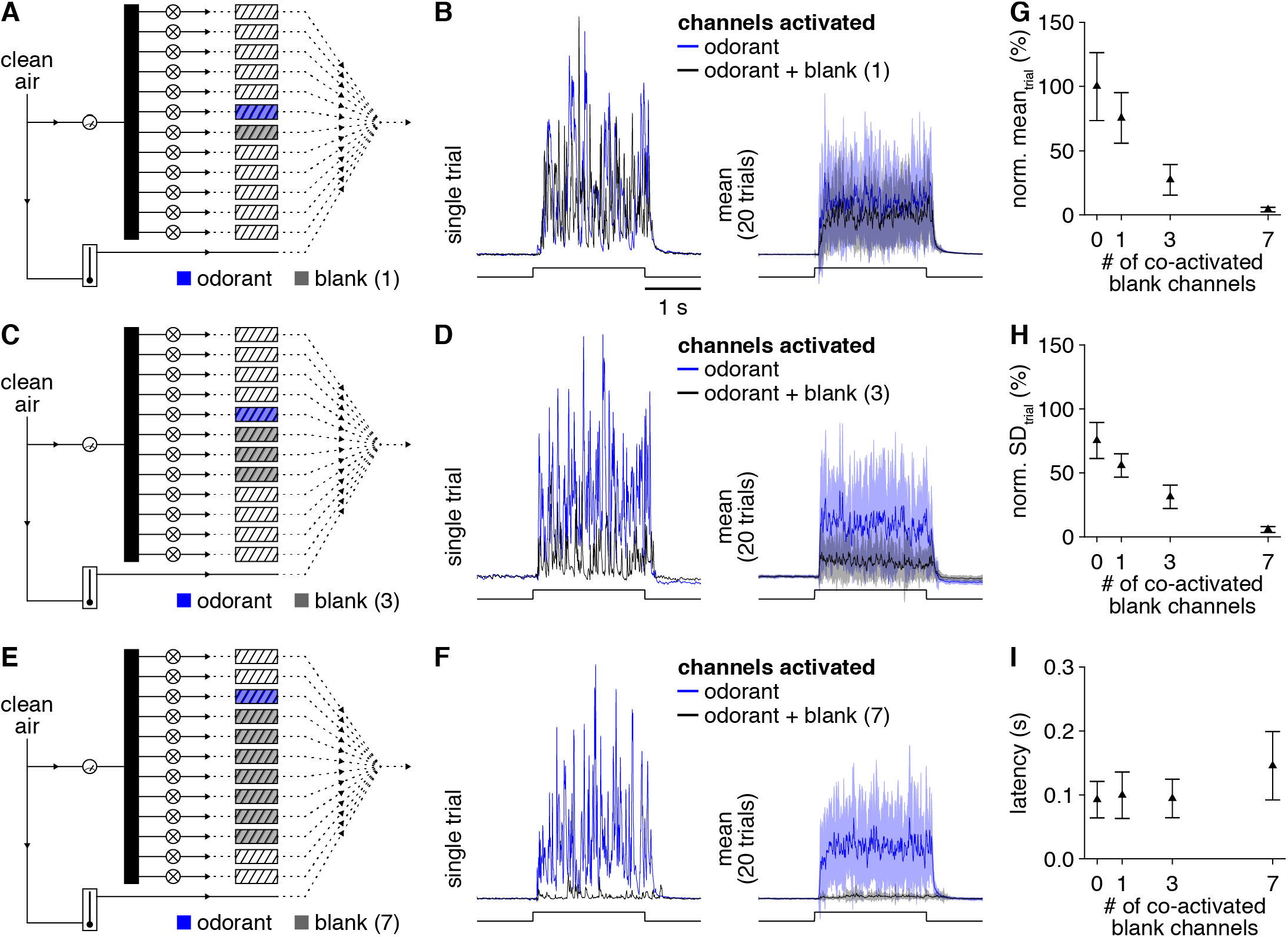
Simple mixture generation with up to 8 individual components. (**A**) Schematic of olfactometer preparation for testing odorant delivery from a single channel in isolation or simultaneously with activation of 1 blank channel (i.e., a channel with non-photoionizable odorant solvent only loaded into the odorant reservoir). (**B**) Single trial PID recording (left) or mean PID recording (right) during 2-s-long valve openings (lower) of the odorant in isolation or the odorant and blank channel simultaneously. (**C-F**) Same as **A,B** for 3 and 7 co-activated blank channels, respectively. (**G,H**) PID recording mean (**G**) and SD (**H**) during the odorant delivery for increasing numbers of co-activated blank channels (normalized to the 0 co-activated blank channels mean). (**I**) Mean latency of odorant delivery for increasing numbers of co-activated blank channels.

### Standard implementation

Based on the above characterization and our experience from daily use of the novel olfactometer, we recommend the following standard practices and operating parameters for use of the device. Odorants are prepared in advance by diluting each to the desired strength in mineral oil or caprylic/capric medium chain triglycerides – a standard food industry solvent (Zhang and Reineccius, 2017) – to enable flexible selection of 12-odorant panels during an experiment. Odorant panels are then generated immediately before or during an experiment by front-loading each odorant reservoir with 40 μL of odorant, which effectively distributes across the internal surface area of the reservoir without leaking from the bottom or impeding airflow. The use of oil as a solvent facilitates this distribution, while water or other hydrophilic solvents do not distribute as well across the plastic substrate of the mixer elements. Including trace quantities of an odorless dye (e.g., Sudan Black B) in the solvent before odorants are prepared further facilitates the loading process by allowing direct visualization of the distributing odorant within the clear-plastic odorant reservoirs. When the experimental preparation is ready, we position the delivery arm of the olfactometer 8-10 cm in front of the preparation, enabling both reliable odorant delivery (**Figure 4**) and easy exchange of the modular barrels without disturbing the preparation. The delivery arm is then locked in place for the duration of the experiment, ensuring equivalent delivery across all odorants tested. A small and quiet air pump with in-line charcoal filter is used to drive carrier stream flow rates of 8 L/min, yielding reliably low-latency odorant delivery (**Figure 6**) as well as a constant delivery of clean air to the experimental preparation between trials. For initial odorant presentation, channel pressures are typically set to 30 kPa. Any desired change in the concentration of odorant delivered is then achieved by modulating the channel pressure by ±10 kPa (for modest changes) or by preparing an odorant reservoir with a different liquid dilution (for large changes).

### *In vivo* validation: neural imaging

Having systematically characterized odorant delivery from the novel olfactometer using PID recordings, we next tested the ability of the olfactometer to drive reliable odorant- and concentration-specific neural activity. Specifically, we compared spatiotemporal patterns of olfactory sensory neuron activation evoked by 3 chemically distinct odorants presented at 2 liquid dilutions using either a flow dilution olfactometer or the novel olfactometer. Consistent with our previous results in anesthetized mice (Wachowiak and Cohen, 2001; Spors et al., 2006), odorant delivery from the flow dilution olfactometer evoked focal glomerular patterns of odorant- and concentration-specific activity in the dorsal main olfactory bulb, with activity temporally modulated to varying degrees by inhalation (**Figure 10A-L**). In turn, turbulent delivery of the same odorants at estimated equivalent mean concentrations from the novel olfactometer evoked essentially identical spatiotemporal activity patterns, with only slight differences in activity likely arising from: 1) imperfectly matched final concentrations, 2) distinct odorant delivery latencies, and/or 3) the slower rise to a final steady-state concentration with the flow dilution olfactometer. Our results thus indicate that odorant delivery from the novel olfactometer evokes reliable odorant- and concentration-specific neural activity comparable to that observed with a flow dilution olfactometer. Moreover, in contrast to flow dilution olfactometers, which can require long intervals for odorant concentrations to equilibrate within tubing and final delivery ports before effective delivery, the novel olfactometer can deliver distinct odorants at arbitrarily short intervals, as demonstrated by the ‘rapid-fire’ presentation of four odorants within 10 seconds (**Figure 10M,N**).

We next turned to a more extensive test of the trial-to-trial reliability and specificity of neural activity evoked by odorant delivery from the novel olfactometer. Capitalizing on the ability of the olfactometer to efficiently deliver numerous odorants, we imaged odorant-evoked activity of olfactory sensory neuron terminals in 105 glomeruli across 73 odorants (including both odorant solvent and empty reservoir negative controls) delivered in a pseudo-random order over 219 trials in a single experimental preparation spanning ~75 minutes from the start of imaging (**Figure 11A**). Across individual trials, odorants reliably evoked distinct glomerular patterns of activity, including activity in just a single glomerulus when particular odorants were presented at low concentration (**Figure 11B**). In contrast, within the span of 219 trials, neither the odorant solvent nor the empty reservoir control evoked any detectable neural activity (**Figure 11C**), confirming a lack of contamination. Finally, correlating responses of all 105 glomeruli across each trial revealed consistently high correlations for repeat trials of the same odorant, while distinct odorants evoked glomerular patterns of activity with variable levels of correlation (**Figure 11D**), as expected from the diverse bank of odorants tested. The novel olfactometer is thus capable of highly efficient and flexible odorant delivery with minimal inter-channel and inter-trial contamination.

**Figure 10.**
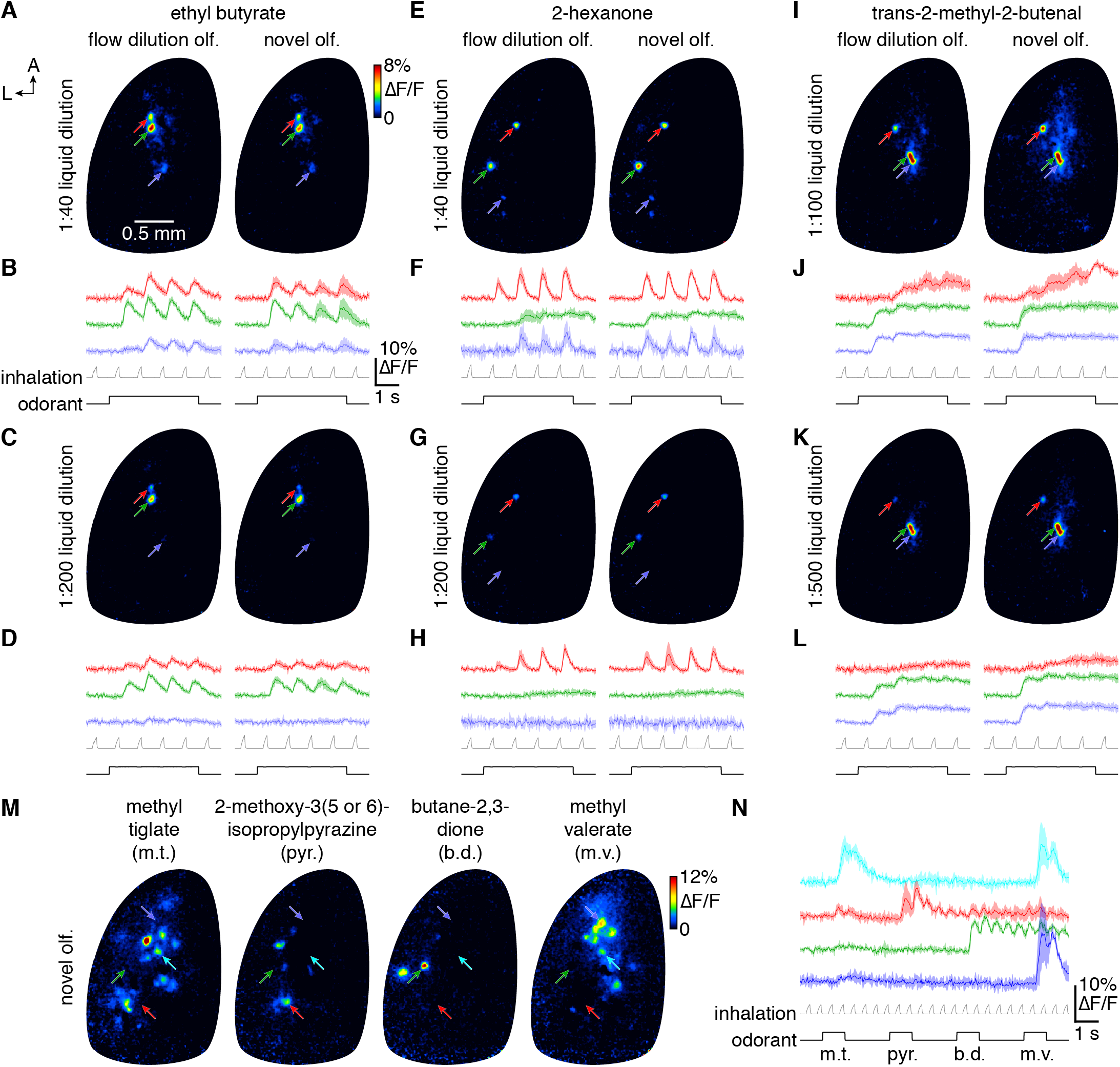
Comparable odorant- and concentration-specific neural activity in the mouse main olfactory bulb evoked by the novel olfactometer and a flow dilution olfactometer. (**A**) Mean ∆F/F map (4 trials) of GCaMP6f responses from olfactory sensory neurons projecting to the dorsal main olfactory bulb of an OMP-Cre;RCL-GCaMP6f mouse following 4-s-presentation of ethyl butyrate (1:40 liquid dilution) by either a standard flow dilution olfactometer (left) or the novel olfactometer (right). (**B**) Mean activity timecourses of the glomerular ROIs labeled in **A** (upper) relative to pressure measurements of artificial inhalation (middle) and the timecourse of odorant presentation (lower). (**C,D**) Same as **A,B** for a lower odorant concentration. (**E-L**) Same as **A-D** for the odorants 2-hexanone (**E-H**) and trans-2-methyl-2-butenal (**I-L**). (**M**) Mean ∆F/F maps (3 trials) following rapid sequential presentation of methyl tiglate, 2-methoxy-3(5 or 6)-isopropylpyrazine, butane-2,3-dione, and methyl valerate. (**N**) Mean activity timecourses of the glomerular ROIs labeled in **M** (upper) relative to pressure measurements of artificial inhalation (middle) and the timecourse of sequential odorant presentation (lower).

**Figure 11.**
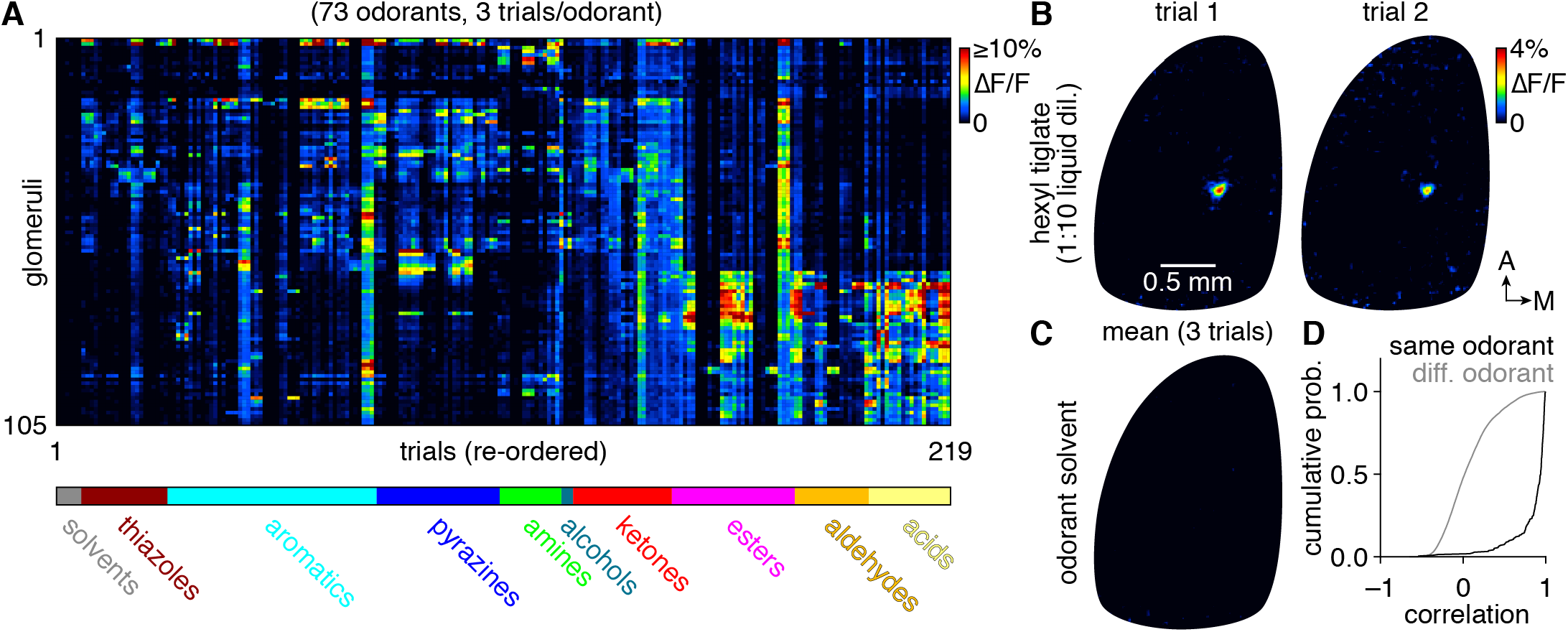
Reliable and efficient odorant delivery. (**A**) Mean ∆F/F GCaMP6f signals from 105 glomeruli in the left main olfactory bulb of an OMP-Cre;RCL-GCaMP6f mouse evoked by 2-s-long delivery of 73 odorants (3 trials/odorant shown; trials re-ordered so that repeat trials of the same odorant appear sequentially). (**B**) Map of ∆F/F signals in the main olfactory bulb summarized in A evoked by hexyl tiglate (1:10 liquid dilution). 2 nonconsecutive trials are shown, with 16 trials delivering 11 other odorants interspersed between the 2 trials. (**C**) Map of ∆F/F signals as in **B**, evoked by the caprylic/capric medium chain triglycerides odorant solvent. Mean of 3 trials shown, collected after delivering 12 chemically diverse odorants, demonstrating no detectable contamination. (**D**) Cumulative probability of inter-trial correlations of glomerular responses shown in **A**, demonstrating consistently high correlations for the same odorants and broadly distributed correlations for different odorants.

### *In vivo* validation: behavior

In our final set of experiments, we investigated the behavioral performance of awake head-fixed mice performing a go/no-go odorant discrimination task using the novel olfactometer. We began by assessing whether mice could discriminate two odorants closely matched in volatility but highly distinct chemically and perceptually: 2-acetylthiazole (CS+) and m-cresol (CS–) (1:10 liquid dilutions). Across three days, mice reached 80% accuracy within the first 100 trials and 90% accuracy within the first 260 trials (**Figure 12A**). While rapid, we note that this rate of learning is slower than some previous studies employing standard flow dilution olfactometers (e.g., (Bodyak and Slotnick, 1999; Chu et al., 2016)), a difference that may reflect our choice of odorants and/or the potentially greater task difficulty imposed by the turbulent odorant delivery.

**Figure 12:**
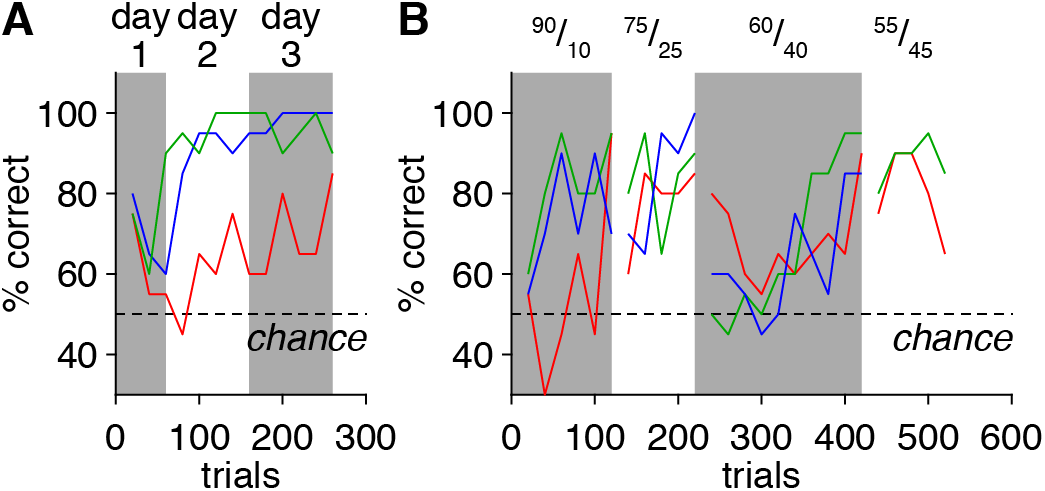
Operant conditioning using the novel olfactometer. (**A**) Behavioral performance of head-fixed mice (n=3) in a simple go/no-go olfactory task requiring discrimination of pure odorants for a water reward. (**B**) Behavioral performance of the same mice in progressively more difficult discrimination tasks requiring discrimination of progressively more similar odorant mixtures.

Following this simple behavioral task, we next evaluated whether the novel olfactometer could be used to administer progressively more difficult tasks by training mice to discriminate binary mixtures of the same two odorants mixed in the liquid phase at varying ratios. Indeed, mice learned to discriminate stimuli ranging from 90% 2-acetylthiazole/10% m-cresol vs. 10% 2-acetylthiazole/90% m-cresol to 55% 2-acetylthiazole/45% m-cresol vs. 45% 2-acetylthiazole/55% m-cresol within 1-2 days, with final performance levels well above chance (**Figure 12B**). Collectively, these results thus demonstrate that the novel olfactometer is well suited for both neural recording and behavioral experiments.

### Alternative designs

The modular and relatively simple design of the novel olfactometer permits a high degree of flexibility in final configuration, depending on the desired performance characteristics. Indeed, not only can numerous operating parameters be easily modulated (e.g., olfactometer-to-experimental preparation distance, channel pressure, carrier stream flow rate, and/or valve opening duration), but multiple components of the olfactometer can also be readily replaced with other commercially available alternatives. For example, mixers for two-part adhesives (i.e., odorant reservoirs) exist in a wide variety of lengths (e.g., 2.9-5.3 in.), permitting different volumes of odorant to be loaded to achieve different degrees of odorant delivery stability. These mixers likewise are available in a variety of tip styles (e.g., blunt, luer slip, luer lock, etc.) that are compatible with different odorant reservoir tips (e.g., dental applicators vs. variable-gauge bent needles), effecting different final bore diameters and consequent odorant delivery rates.

Beyond replacing individual components, the novel olfactometer design can also be easily added to, as demonstrated by the addition of an inexpensive mixing eductor (**Table 1**) to the carrier stream tube (**Figure 13A**) to enhance mixing of the odorant within the carrier stream and reduce variance in the instantaneous concentration of odorant delivery. Briefly, an eductor combines an upstream nozzle and downstream conical diffuser separated by a short open span. As fluid (vapor or liquid) passes through the constricting nozzle its velocity increases, generating a decrease in pressure (i.e., the Venturi effect) that draws in surrounding fluid through the open span. The central stream and drawn-in fluid then enter the diffuser at high velocity, whereupon the broadening walls generate turbulent mixing, ultimately outputting a well-mixed stream.

To implement this modification, we attach the eductor to the carrier stream tube and aim the odorant reservoir tips toward the open span of the eductor, allowing expelled saturated odorant vapor (plus surrounding air) to be drawn into the diffuser (**Figure 13B**) and thoroughly mixed with the carrier stream (**Figure 13C**). PID measurements of odorant delivery (**Figure 13D,E**) revealed that addition of the eductor reduced the variance in the instantaneous concentration of odorant delivery (i.e., PID signal SD) by ~76%, which equated to a ~32% reduction in the PID signal coefficient of variation during the odorant pulse (**Figure 13G**) when accounting for the ~66% increase in mean odorant dilution (**Figure 13F**). Addition of the eductor also dramatically reduced the variability in latency from valve opening to odorant delivery by ~82% (**Figure 13D**).

Three caveats must be considered with the optional addition of an eductor. First, the struts connecting the eductor nozzle and diffuser can physically occlude odorant delivery from a subset of the 12 total olfactometer channels, necessitating greater care in positioning of odorant reservoir tips. Second, odorant passing through the eductor is mixed with air drawn in from the surrounding environment, increasing the potential for nonspecific odorant input to the experimental preparation. Exhaust systems positioned around the experimental station can mitigate this effect, however. Finally, the eductor introduces an adsorptive surface to the novel olfactometer that necessarily increases the probability of inter-trial contamination. We note, however, that spans of open airflow are still maintained both upstream and downstream of odorant reservoirs independent of eductor addition. Moreover, the relatively laminar flow within the upstream portion of the eductor preserves a boundary layer between incoming saturated odorant vapor and the eductor body, minimizing odorant adsorption. Finally, the modular design of the olfactometer allows for an eductor to be quickly removed and replaced with one of multiple clean eductors at hand during the course of an experiment, further minimizing concerns of inter-trial contamination.

**Figure 13.**
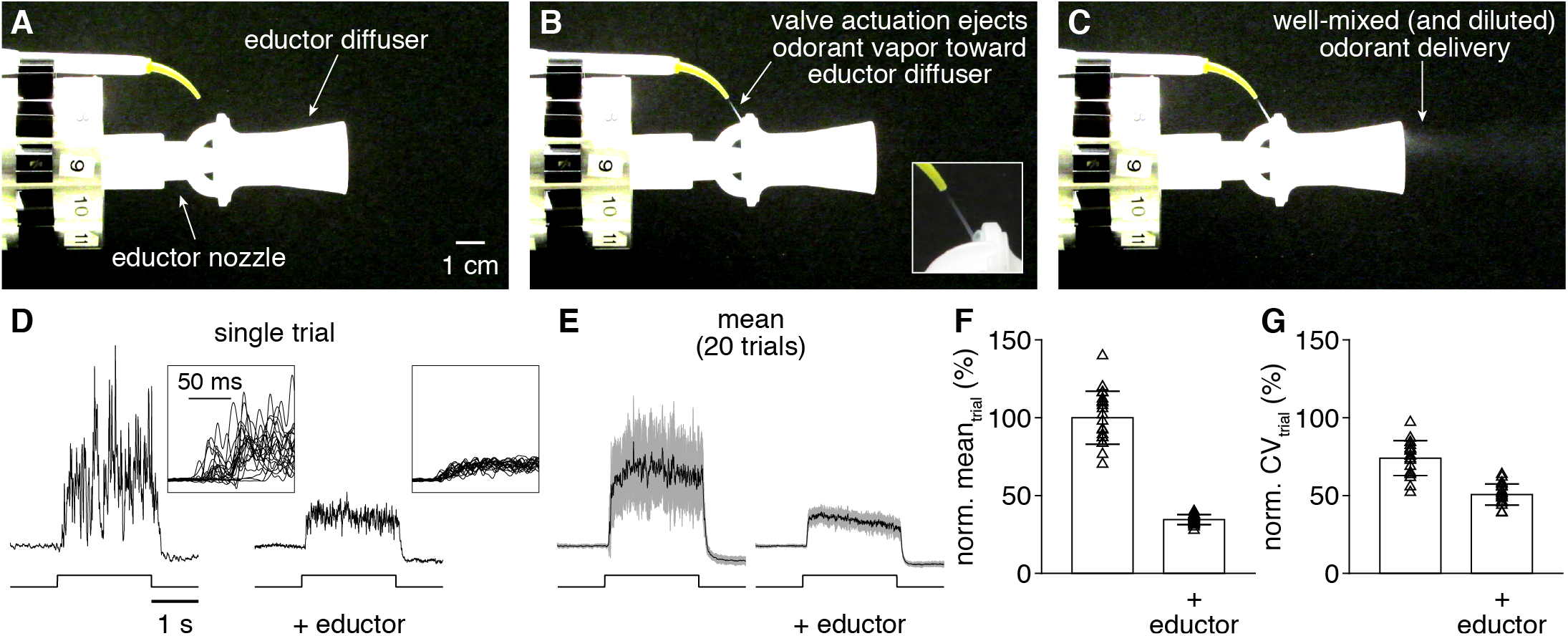
Optional addition of an educator can reduce odorant delivery variability. (**A-C**) Images of TiCl_4_ delivery from the novel olfactometer with eductor immediately before valve actuation (**A**), immediately following valve actuation (**B**; inset: magnification of expelled saturated odorant vapor being drawn into eductor diffuser), and following mixing of the expelled TiCl4 with the carrier stream (**C**). (**D**) PID recordings (upper) during single trials of 2-s odorant delivery (lower) from a single olfactometer channel without or with eductor addition. Inset: magnification of odorant delivery onset across 20 trials, demonstrating the reduction in variability of odorant delivery latency with eductor addition. (**E**) Mean PID recording across 20 trials without or with eductor addition. (**F,G**) PID recording mean (**F**) and CV (**G**) during odorant delivery without or with eductor addition (normalized to the no-eductor mean).

## DISCUSSION

We have presented a novel olfactometer design and validated its performance using PID recordings, neural imaging, and behavioral testing. The distinguishing novel features of the olfactometer are its mixing of saturated odorant vapor with a carrier stream in free space, and the isolation of odorant reservoirs from the delivery flow path by spans of open airflow. These features greatly reduce the available surfaces to which odorants can adsorb as well as the possibility of odorant backflow into upstream manifolds, which collectively minimize both inter-trial and inter-channel contamination of one odorant with another. Although resulting in turbulent odorant delivery, this design strategy nevertheless permits easy experimental control over the mean concentration, onset latency, and duration of odorant delivery, and further yields delivery that is not only indistinguishable across 12 independent channels but also reliable across trials. Indeed, the trial-to-trial reliability and consistency of odorant delivery enabled different odorants and concentrations to be mapped to neural activity with a level of precision comparable to that obtained with a flow dilution olfactometer, and further supported operant conditioning of mice in an odorant discrimination task. Finally, from a practical standpoint, the novel olfactometer is easily assembled from mostly commercially available components, is relatively inexpensive to construct and use, and its simple design allows for a range of modifications to adapt the device to particular experimental applications.

### Advantages of the novel olfactometer

A primary motivation for developing this novel design was the need to efficiently and flexibly test odorants in the context of mapping sensory information to neural activity in the olfactory system. Addressing this problem requires testing responses to large numbers of odorants – often at multiple concentrations – due to the: 1) high dimensionality of olfactory stimulus space, 2) limited knowledge of odorant receptor tuning, and 3) limited knowledge of (and animal-to-animal variability in) glomerular positions. Several previous studies have approached this problem by constructing flow dilution olfactometers capable of delivering relatively large numbers of odorants (64-100 per experiment) (e.g., see: (Davison and Katz, 2007; Soucy et al., 2009; Tan et al., 2010)) and their results have provided important insights into olfactory processing. However, with each of these olfactometers, the control of odorant delivery with dedicated lines requires odorant panels to be pre-selected. The requirement to replace these lines and, in some cases, control valves when introducing a new odorant or a new concentration range creates an energy barrier to expanding or modifying a chosen odorant panel. In addition, the use of a common final delivery line, as well as a common upstream supply line, introduces substantial possibility for inter-trial and inter-channel contamination. In our experience with simpler versions of such flow dilution olfactometers, we have found these issues to pose serious bottlenecks to comprehensively testing large odorant panels, with issues such as inter-trial and inter-channel contamination becoming increasingly apparent with recent improvements in the sensitivity of tools for monitoring odorant-evoked neural activity.

The novel features of the olfactometer presented here largely solve the issues described above. These features, together with the modular design and integration of disposable odorant reservoirs, enable rapid exchange of 12-odorant panels during experimentation, supporting highly efficient and flexible odorant testing. With such a design, odorants and concentrations can be chosen even during the course of an experiment, and the number of total stimuli that can be tested is limited only by the endurance of the experimental preparation and experimenter.

Several additional advantages also emerged as a result of the novel design. First, because each olfactometer channel is essentially independent, delivering mixtures of nearly any combination of 12 odorants in vapor phase is trivial. Although such mixture generation is somewhat limited in the current implementation by the division of airflow over simultaneously open valves, resulting in a drop in channel pressure (and thus mean concentration of odorant delivery) with increasing mixture complexity, this issue could easily be resolved by using a separately regulated pressure source for each channel. Second, unlike flow dilution olfactometers, there is no time required for washout of odorant from a common final delivery line or for an odorant to reach steady-state concentrations within a mixing chamber. As a result, there is practically no limitation to the timing of odorant delivery from different channels: different odorants can be delivered at arbitrarily short intervals relative to one another, and can even overlap in time. This temporal flexibility dramatically increases the efficiency with which odorants can be tested, and may further prove useful in examining temporal relationships in olfactory processing. Finally, the novel olfactometer may be of particular utility for investigating the neural encoding and behavioral perception of naturalistically fluctuating odorant stimuli. Such an application is especially well suited for non-mammalian model organisms in which rapidly fluctuating odorant stimuli profoundly shape neural activity, such as locusts (e.g., see: (Geffen et al., 2009; Aldworth and Stopfer, 2015; Huston et al., 2015)), moths (e.g., see: (Vickers et al., 2001)), and flies (e.g., see: (Nagel and Wilson, 2011; Nagel et al., 2014)).

### Limitations of the novel olfactometer

Consistent with any olfactometer operating with liquid odorant dilutions, final absolute odorant concentrations delivered from our novel olfactometer are difficult to accurately predict due to non-ideal behavior of odorants at the liquid-to-vapor transition (Slotnick and Restrepo, 2005). Incomplete mixing of expelled saturated odorant vapor with the carrier stream in the novel olfactometer design (**Figure 6**) exacerbates this difficulty. Thus, the novel olfactometer is not ideal for absolute measurements of concentration-response functions in terms of odorant molarity. Nevertheless, the novel olfactometer can effectively deliver distinct relative concentrations, both through easily prepared liquid dilutions that are loaded into the disposable odorant reservoirs, as well as through instantaneous changes in channel pressure (**Figure 5**). We have found this level of control to easily support rapid testing of odorants across a wide concentration range, including exceedingly low concentrations capable of revealing highly selective activation of olfactory sensory neurons projecting to single glomeruli.

The novel olfactometer can effectively deliver odorants across a large range of volatility (**Figure 8; Table 2**). However, due to the open nature of the odorant reservoirs, with no immediately upstream or downstream tubing, extremely volatile odorants (e.g., N,N-dimethylethylamine, vapor pressure: 495.357 mmHg) can escape the odorant reservoir and enter the carrier stream even when the corresponding channel valve is not actuated (data not shown). The novel olfactometer is thus not suitable for testing such extremely volatile odorants without further modification. While we have not systematically examined over what range of volatility the novel olfactometer can achieve tightly controlled odorant delivery, in practice we have had excellent success monitoring neural activity in the mouse olfactory system time-locked to channel valve actuation with the vast majority of odorants tested.

Olfactory information is encoded in precise spatiotemporal patterns of neural activity within both vertebrate and invertebrate olfactory systems (Laurent et al., 2001; Schaefer and Margrie, 2007; Chong and Rinberg, 2018). Turbulent odorant delivery from the novel olfactometer may complicate the investigation of such dynamic neural activity by introducing additional temporal variance. Placement of a PID close to the experimental preparation to simultaneously monitor odorant delivery and neural activity could help account for this added complexity. Moreover, variance in the instantaneous concentration of odorant delivery can be substantially reduced by the addition of an eductor to achieve more thorough mixing of the expelled saturated odorant vapor with the carrier stream (**Figure 13**). At the same time, inhalation (in mammalian preparations) may mitigate this potential confound by low-pass filtering turbulent stimuli. Indeed, using ultrasensitive GCaMP6f to monitor olfactory sensory neuron activity, we observed no difference in the variability of neural activity evoked by turbulent odorant delivery from the novel olfactometer compared to that evoked by comparatively stable odorant delivery from a standard flow dilution olfactometer (**Figure 10**). Whether comparable patterns of neural activity are maintained using measures of activity with greater temporal resolution than GCaMP6f (e.g., electrophysiological recordings) and in awake, behaving mice warrants further investigation.

## ACKNOWLEDGMENTS

We thank Jackson Ball, Gustavo A. Vasquez-Opazo, Thomas Rust, and Rebecca L. Kummer for excellent technical assistance and Daniel W. Wesson, Andrew K. Moran, Shaina M. Short, and Isaac A. Youngstrom for helpful discussion.

